# A Framework for Quantifying DNA Methylation Heterogeneity and Detecting Co-methylated loci from Native Nanopore Sequencing

**DOI:** 10.64898/2026.09.07.749820

**Authors:** Young Jun Kim, Nicolae Radu Zabet

## Abstract

DNA methylation is an important epigenetic mechanism involved in gene regulation. Most methods focus on analysing DNA methylation averaged from multiple reads, yet these average methylation profiles obscure heterogeneity between individual DNA molecules and coordinated methylation states across loci. Native Oxford Nanopore Technology (ONT) sequencing directly captures long, native DNA molecules together with their base modifications, allowing methylation to be studied at single-molecule resolution. Here, we present a scalable framework for genome-wide methylation analysis using ONT sequencing at single-molecule resolution, focused on two features that site-level summaries cannot recover. First, it quantifies molecule-to-molecule heterogeneity in DNA methylation by detecting Variable Methylated Domains (VMDs) and Variable Methylated Regions (VMRs). Second, it identifies coordinated methylation, as co-methylated positions (CMPs) and regions (CMRs), from the states observed on the same individual DNA molecules. Using data from human LCL cells, we demonstrate that substantial molecule-level methylation heterogeneity is masked by site-level summaries. Co-methylation analysis reveals coordinated patterns between CpG sites and genomic regions, including shared and sex-specific patterns, uncovering methylation organisation not apparent from average methylation levels. We also show that CMPs can be used to detect TF-pairs that are predicted to have coordinated binding. This framework is integrated within the DMRcaller R/Bioconductor package.

## Introduction

DNA methylation (DNAm) is an epigenetic modification involving the addition of a methyl group to cytosine residues, predominantly at CpG dinucleotides. This evolutionarily conserved process plays a fundamental role in chromatin organisation, gene regulation, and genomic stability (1; 2), and has significant clinical relevance in cancer as both a diagnostic and mechanistic biomarker (3). Consequently, genome-wide analysis of DNA methylation patterns is essential for understanding epigenetic regulation in development and disease.

Given the reduction in cost of DNA sequencing, Whole Genome Bisulfite Sequencing (WGBS-seq) has become the gold standard in the field for building genome-wide DNAm maps at base pair resolution. In WGBS-seq, the bisulfite treatment of the DNA leads to unmethylated cytosines being converted to uracil and 5-methylcytosines remaining unaffected. The process also includes a PCR amplification which converts uracil to thymine. The bisulfite treatment causes significant DNA degradation (4). Recent advances in long-read sequencing technologies, particularly Oxford Nanopore Technologies (ONT), have enabled direct detection of DNA methylation from native DNA molecules without bisulfite conversion (5; 6). By preserving long-range genomic context, ONT data provide unique opportunities to investigate methylation heterogeneity, long-range epigenetic structure, and coordinated methylation patterns across individual DNA molecules. This single-molecule resolution offers a substantial advantage over short-read sequencing approaches, which rely on population-level averaging and often fail to capture epigenetic variability across cells or alleles (7).

To leverage these capabilities, several computational tools have been developed to support methylation analysis from ONT data, including methods for base-level modification calling such as Nanopolish, Megalodon, and Dorado, as well as tools for regional aggregation such as Modkit (5). Although ONT-based methods enable direct methylation calling at the single-molecule level, most existing computational workflows primarily report site-level or region-level summaries rather than detailed genome-wide single-molecule methylation patterns (8; 9). Almost all downstream tools reduce the data to per-site methylation: the reads covering a cytosine are pooled into a single value (for ONT this is usually done with Modkit) that is then passed to packages developed for WGBS, such as methylKit (33) or DSS (34). Once the reads have been pooled in this way, the information about which CpGs are methylated together on the same molecule is lost and cannot be reconstructed from the per-site values. This means that important features such as co-methylation structure, long-range coordination, and regional methylation heterogeneity remain insufficiently explored.

Of these underexplored features, heterogeneous DNA methylation patterns are of particular biological and clinical significance, as intercellular epigenetic diversity has been linked to key processes in development, ageing, and disease (10; 11). Partially methylated domains (PMDs), which represent extended genomic regions of intermediate methylation, have emerged as hallmarks of epigenetic heterogeneity in specific biological contexts, including cancer cells, the placenta, and cultured cell lines (12; 13). Within tumours, methylation heterogeneity acts as an independent driver of clonal evolution, metastasis, and resistance to therapy, functioning as a non-genetic mechanism of cancer progression that is distinct from, yet complementary to, somatic mutation (14; 15). Beyond PMDs, variable methylated domains (VMDs) provide a complementary framework for quantifying the degree of read-level stochasticity within partially methylated regions, offering a finer-grained view of epigenetic heterogeneity that is not accessible through population-averaged methylation summaries (7). Taken together, the systematic characterisation of heterogeneous methylation patterns represents a critical step towards understanding the epigenetic underpinnings of both normal cellular diversity and disease progression.

Here, we present a new computational framework that enables genome-wide methylation analysis from native ONT sequencing, supporting the identification of molecule-level DNA methylation heterogeneity, and co-methylation patterns directly from ONT-derived methylation calls. By integrating long-read methylation information within a unified statistical framework, *DMRcaller* provides a flexible and scalable platform for exploring epigenetic variation across genomes using Nanopore sequencing data, filling a critical gap in the current landscape for single-molecule methylation analysis.

## Materials and Methods

### Cell lines and sequencing data

All analyses were performed using publicly available Oxford Nanopore Technologies (ONT) sequencing data from the 1000 Genomes ONT dataset (17), which provides whole-genome long-read sequencing data for a diverse panel of human lymphoblastoid cell lines (LCLs). Three cell lines were used in this study: GM18501 (male, Yoruba ancestry) for single-sample analysis, and HG00277 (male, Finnish ancestry) and HG00326 (female, Finnish ancestry) for sex-differential methylation analyses. Raw signal data in .pod5 format were obtained directly from the 1000 Genomes ONT public S3 repository (https://s3.amazonaws.com/1000g-ont/).

### DNA methylation analysis for ONT data

Figure 1 represents an overview of the workflow for DNA methylation analysis with DMRcaller. Individual steps are detailed below.

**Figure 1:**
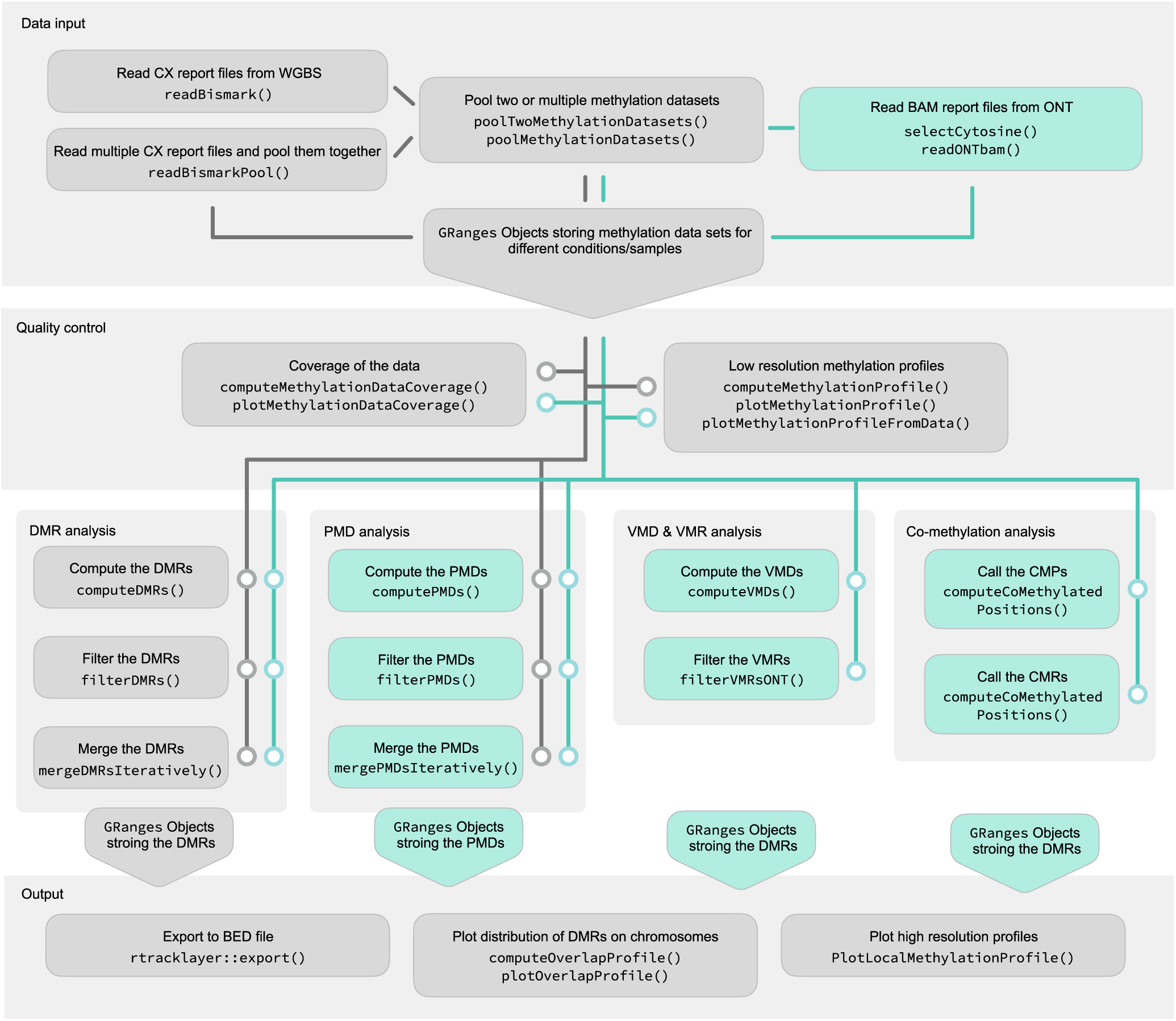
Overview of the DMRcaller analytical framework for long-read methylation analysis. Schematic representation of the new analytical framework introduced in *DMRcaller*, illustrating the progression from population-level differential methylation (DMRs with computeDMRs, filterDMRs and mergeDMRsInteratively; and PMDs with `computePMDs`, `filterPMDs` and `mergePMDsInteratively`) to single-molecule heterogeneity (VMDs with `computeVMDs` and VMRs with `computeVMRs`) and coordinated methylation structure (CMPs with `computeCoMethylatedPositions` and CMRs with `computeCoMethylatedRegions`) at individual genomic loci. Green colours steps highlights the new added functionality for ONT DNAm data. The ONT DNAm data is imported using `readONTbam` function.

#### Basecalling and alignment

Basecalling, base modification calling, and alignment were performed simultaneously using Dorado v0.9.6 (Oxford Nanopore Technologies). For the GM18501 validation cell line, which was sequenced on an R9.4.1 flow cell, basecalling was performed using the high-accuracy (`hac`) model with CpG 5mC modification calling (dna_r9.4.1_e8_hac@v3.3 with `5mCG` modification). For `HG00277` and `HG00326`, used for sex-differential methylation analyses, the dna_r10.4.1_e8.2_400bps_sup@v5.2.0 modification model was applied with Dorado v1.1.0, enabling joint detection of 5-methylcytosine (5mC) and 5-hydroxymethylcytosine (5hmC) alongside high-accuracy basecalling. Reads were aligned to either the GRCh38 (hg38) reference genome or the `T2T-CHM13v2.0` reference genome (GenBank accession: `GCA_009914755.4`) (18) depending on the analysis. For sex-differential methylation analyses, T2T-CHM13v2.0 was used as the primary reference to improve alignment accuracy in centromeric and pericentromeric regions, which are composed of 171-bp *α*-satellite monomers organised into higher-order repeats that are poorly resolved in `GRCh38` (19; 20). A minimum base quality score threshold of 10 was applied during basecalling. The resulting BAM files contain `MM` (modified base) and `ML` (modification probability) tags encoding per-read base modification calls at single-molecule resolution.

#### Methylation extraction using *DMRcaller*

Per-cytosine methylation statistics were extracted from the aligned BAM files using the new `readONTbam` function in *DMRcaller* package. This function decodes `MM/ML` tags and computes, for each CpG site in the reference genome, the number of methylated reads (`readsM`), total reads (`readsN`), and per-read methylation indices, storing the output as a `GRanges` object. Cytosine positions in CG context were identified from the T2T-CHM13v2.0 reference genome(18) using the `selectCytosine` function with the `BSgenome.Hsapiens.NCBI.T2T.CHM13v2.0` Bioconductor package. The `readONTbam` function was run per chromosome with a probability threshold of 0.5, above which a modification call was considered methylated.

#### Global methylation profiling

Low-resolution global methylation profiles were generated using the `computeMethylationProfile` function with non-overlapping 1 Mb windows across all chromosomes. Mean methylation proportion and mean sequencing coverage were computed per window in CG context and visualised using `plotMethylationProfileFromData`. Profiles were generated separately for each cell line to enable visual comparison of global methylation levels and coverage distributions.

#### Partially Methylated Domains detection

Partially methylated domains (PMDs) were identified using the new `computePMDs` function in *DMRcaller*. For single-sample validation (GM18501), PMDs were defined as domains with mean methylation proportions between 0.4 and 0.6. Three detection methods were applied and compared: noise-filter (with a triangular kernel, *λ* = 0.5, window size 100 *bp*), neighbourhood, and bins (bin size 100 *bp*). Common parameters across all three methods were: `minCytosinesCount` = 4, `minGap` = 200 bp, `minSize` = 200 bp, and `minReadsPerCytosine` = 4. For sex-differential co-methylation analyses (HG00277 and HG00326), PMDs were identified using the Bins method with an expanded methylation proportion range of [0.2 − 0.8] and a minimum domain size (`minSize`) of 1 *Kb*, to capture a broader set of partially methylated regions suitable for co-methylation profiling.

#### Variable Methylated Domains detection

Variable methylated domains (VMDs) were identified using the new computeVMDs function. For GM18501, the genome was segmented into fixed-size bins ( binSize = 200 bp), and the weighted standard deviation (SD) of per-read methylation proportions was computed within each bin. High- and low-variability VMDs were defined using the EDE.high and EDE.low methods respectively, which apply the *ede* elbow-detection algorithm from the R package *inflection* to the empirical SD distribution to identify knee points separating high-variance from low-variance bins. VMD detection was restricted to the PMD search space identified in the GM18501 cell line. Parameters used were: minCytosinesCount = 4, minGap = 200 bp, minSize = 50 bp, and minReadsPerCytosine = 4.

For sex-differential methylation analyses (HG00277 and HG00326) filterVMDs was applied to the common PMD set (the 152), and, separately, the promoter regions (2 *Kb* upstream of TSS) using minCytosinesCount = 4, minReadsPerCytosine = 10; and selecting the high/low VMDs (top/bottom 10% each sex).

#### Variable Methylated Regions detection

Variable methylated regions (VMRs) between HG00277 and HG00326 were identified using the `filterVMRsONT` function, applied at PMDs common between males and females and at promoters (2 kb upstream of TSS) in the T2T-CHM13v2.0 genome annotation (GenBank accession: GCA_009914755.4) (18). The function applies two statistical tests to each candidate region: a Mann-Whitney U test on per-read methylation proportions to assess differences in central tendency, and an F-test on variance of per-read methylation proportions to assess differences in dispersion. P-values from both tests were adjusted for multiple testing using the Benjamini–Hochberg method (21). Parameters used were: `pValueThreshold` = 0.05, `minCytosinesCount` = 4, `minProportionDifference` = 0.4, `minReadsPerCytosine` = 10, and `ciExcludesOne = TRUE`. Regions identified as both differentially methylated (is_DMR = TRUE) and variable methylated (`is_VMR = TRUE`) were retained as DMR–VMR co-localising regions.

#### Differentially Methylated Regions detection

Sex-associated differentially methylated regions (saDMRs) between HG00277 and HG00326 were identified using the `computeDMRs` function with the bins method in *DMRcaller* (16). DMR detection was performed genome-wide in CG context using the bins method with the following parameters: `test = “score“`, `pValueThreshold` = 0.05, `minCytosinesCount` = 1, `minGap` = 200 bp, `minSize` = 50 bp, and `minReadsPerCytosine` = 2.

In order to detect DMRs, we selected the bins method with a bin size of 200 bp and then tested several methylation proportion difference thresholds ranging from 0.05 to 0.9 to determine an optimal threshold (Figure S1A). As the threshold increased, the number of detected saDMRs declined approximately linearly, reflecting the expected trade-off between stringency and sensitivity.

Finally, to determine the optimal bin size, we examined the effect of varying bin size on saDMR genomic coverage on chromosome 1 (Figure S1B). The total size of detected saDMRs declined negligible as bin size increased up to 400 bp, beyond which there was a marked reduction. Altogether, we selected the bins method with a bin size of 400 bp and a min proportion difference of 0.4.

#### Co-methylation analysis

Co-methylated positions (CMPs) were first identified using the `computeCoMethylatedPositions` function to assess pairwise methylation concordance between CpG sites within individual reads. For each pair of CpG sites within a given region, a 2*×*2 contingency table was constructed from read-level co-occurrence counts (both methylated, first only, second only, or neither), and statistical significance was assessed using Fisher’s exact test. Three alternative hypotheses were evaluated and compared: two-sided (detecting any significant association), `greater` (detecting positive co-methylation, where both CpGs are preferentially methylated or unmethylated on the same molecule), and less (detecting anti-co-methylation, where the methylation states at the two CpG sites are inversely correlated). For the GM18501 data, we used the following parameters to identify CMPs: `minDistance` = 150 bp, `maxDistance` = 1,000 bp, `minCoverage` = 1, and `pValueThreshold` = 0.01. For HG00277 vs HG00326, the parameters for CMP detection were: `minDistance` = 0 bp, `maxDistance` = 2,000 bp, `minCoverage` = 4, and `pValueThreshold` = 0.05.

Co-methylated regions (CMRs) were identified using the `computeCoMethylatedRegions` function. For each pair of genomic regions within a user-defined distance range, co-methylation was assessed by computing the Pearson correlation coefficient between the average CpG methylation proportions of the two regions across individual reads. P-values were adjusted using the Benjamini–Hochberg method (21). Parameters used were: `minDistance` = 500 bp, `maxDistance` = 50,000 bp, `minCoverage` = 4, `minCorrelation` = −0.5, `maxCorrelation` = 0.5, and `pValueThreshold` = 0.05. For the sex differences analysis, CMPs were considered common if the overlap between the positions of the two CpGs is exactly matching between males and females, while, CMRs were considered common if the overlap between the two loops is at least 80%.

#### TF co-binding prediction from CMP data

To identify pairs of TFs that co-bind, we selected all TF motifs that have a CpG (with a minimum value for the corresponding base of 0.5 within the PWM motif), resulting in 509 human TF motifs from PWMEnrich package (22). Next, from the 509 TFs with a CpG site in the motif, we selected only TFs that are expressed (with a TPM of at least 1) in a representative LCL line (GM12878) (23), which resulted in 305 TFs. We then performed TF motif enrichment analysis for these 305 TFs using PWMEnrich package (22) and found 27 TFs with motifs enriched at the CMPs in GM18501 cell line. For each of these TFs we computed their predicted binding sites based on PWM score using PWMEnrich package (22) and selecting a minimum raw score of 5. Finally, we counted how many CMPs link binding sites for pairs of TFs (the CpG from the CMP needed to overlap the TF binding site). We performed this analysis only on CMPs that connect CpGs within 150 bp of each other (64,630) in GM18501 cell line.

#### Non-CpG and 5hmC methylation profiling

To demonstrate the multi-context capability of *DMRcaller*, HG00277 ONT sequencing data were processed using Dorado v0.9.6 with the `sup,5mC_5hmC` model to extract base modification calls in six cytosine contexts: CG_5mC, CHG_5mC, CHH_5mC, CG_5hmC, CHG_5hmC, and CHH_5hmC. The `readONTbam` function was applied with the corresponding modif parameter for each context (`“C+m?“` for 5mC and `“C+h?“` for 5hmC). Global methylation proportion and coverage profiles were generated using 1 Mb non-overlapping windows across chromosome 22 for each modification context.

#### Genomic annotation

saDMRs and saDMPs from (24) were annotated against the T2T-CHM13v2.0 reference genome annotation (GenBank accession: GCA_009914755.4) (18), assigning each element to one of eight genomic feature categories: intergenic regions, promoters (defined as 2 kb upstream of TSS), introns, exons, 5’ UTRs, 3’ UTRs, intronic transposable elements (TEs), and intergenic TEs. The proportion of saDMRs and saDMPs overlapping each feature category was compared against the whole-genome baseline proportion of each feature.

#### Statistical analyses and software

All analyses were performed in R v4.5.1 using the updated *DMRcaller* package (16), with parallelisation via the *BiocParallel* package (25). Genome coordinate operations were performed using *GenomicRanges* (26). Visualisation of co-methylation patterns and sequencing data was performed in IGV v2.19.4 (27; 28). Mann-Whitney U tests and Fligner–Killeen tests for comparison of VMD distributions were performed using base R stats functions. All p-values from multiple comparisons were adjusted using the Benjamini–Hochberg false discovery rate procedure(21).

The updated DMRcaller package is available on Bioconductor(29; 30) at https://bioconductor.org/packages/ DMRcaller/. The scripts to perform the analysis presented in this manuscript can be found on Zenodo (31).

## Results

Whole Genome Bisulfite Sequencing (WGBS-seq) is the current gold standard method to generate genome-wide DNAm maps at base pair resolution. The common approach includes pooling all reads over a cytosine and computing the percentage of methylated reads at that position. This is usually done by tools such as Bismark (32) and the data is then input in R/Bioconductor packages that can compute differential methylation, such as MethylKit (33), DSS (34) or DMRcaller (16); see Figure 2. Recently, Oxford Nanopore Technologies (ONT) sequencing has enabled direct detection of DNA methylation from native DNA molecules without bisulfite conversion and PCR amplification (5; 6). Modkit (5) can compute the percentage methylation for each position in the genome, which allows downstream integration of the data with all tools developed for WGBS-seq. Nevertheless, by doing so, information on the actual DNA molecule is lost and analysis investigating methylation at single DNA molecule is not possible. We have now developed a new framework for direct reading of ONT DNA methylation with maintenance of information on the read where the methylation state was measured that is now integrated in DMRcaller (16); see Figure 2.

**Figure 2:**
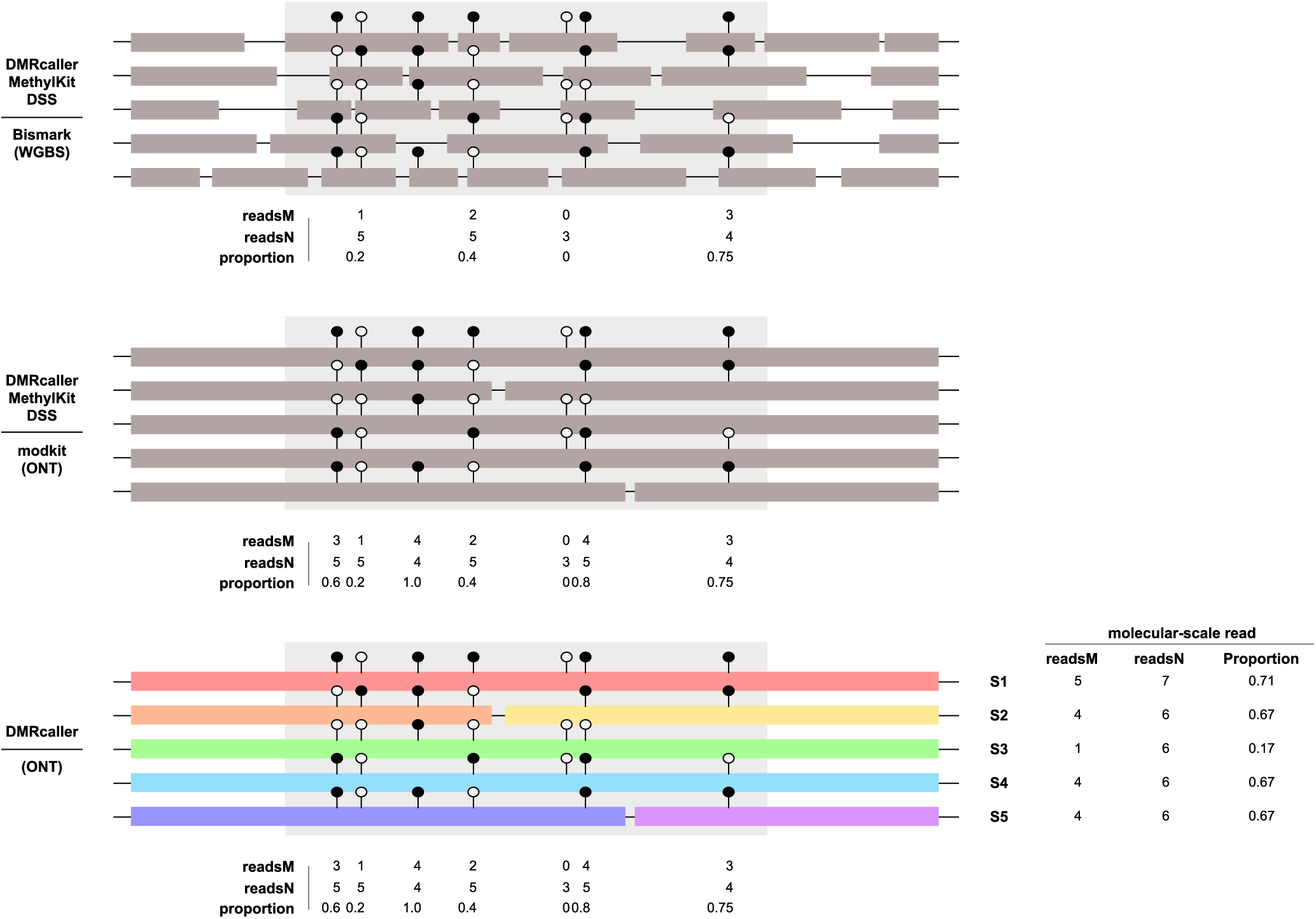
Comparison between WGBS and ONT methods for bulk and single molecule analysis. Schematic overview of the main differences between *(top)* WGBS treated as bulk DNAm (Bismark (32)) and using MethylKit (33), DSS (35) or DMRcaller (16); *(middle)* ONT treated as bulk DNAm (ModKit (5)) and using MethylKit, DSS or DMRcaller; and *(bottom)* ONT treated as single molecule DNAm (*DMRcaller*) and using DMRcaller. *DMRcaller* is reading directly the Dorado-based basecalling and base modification calling bam files using `readONTbam` function. The output `GRanges` object containing single-molecule methylation indices serves as input for downstream analyses including PMD, VMD, VMR, CMP, CMR, and DMR calling.

### Analysing co-methylation and variability in DNA methylation in single samples

To evaluate the new framework for analysing Nanopore sequencing data, we used the GM18501 human lymphoblastoid cell line (LCL) from the 1000 Genomes ONT dataset(17). The average coverage of GM18501 is approximately 15 reads across most regions, except for centromeric and pericentromeric regions, while 5mC proportion is generally between 0.6 and 0.8 (Figure 3A-B and Figure S2A). Note that ONT distinguishes between 5mC and 5hmC and in this analysis, we focused on 5mC only.

**Figure 3:**
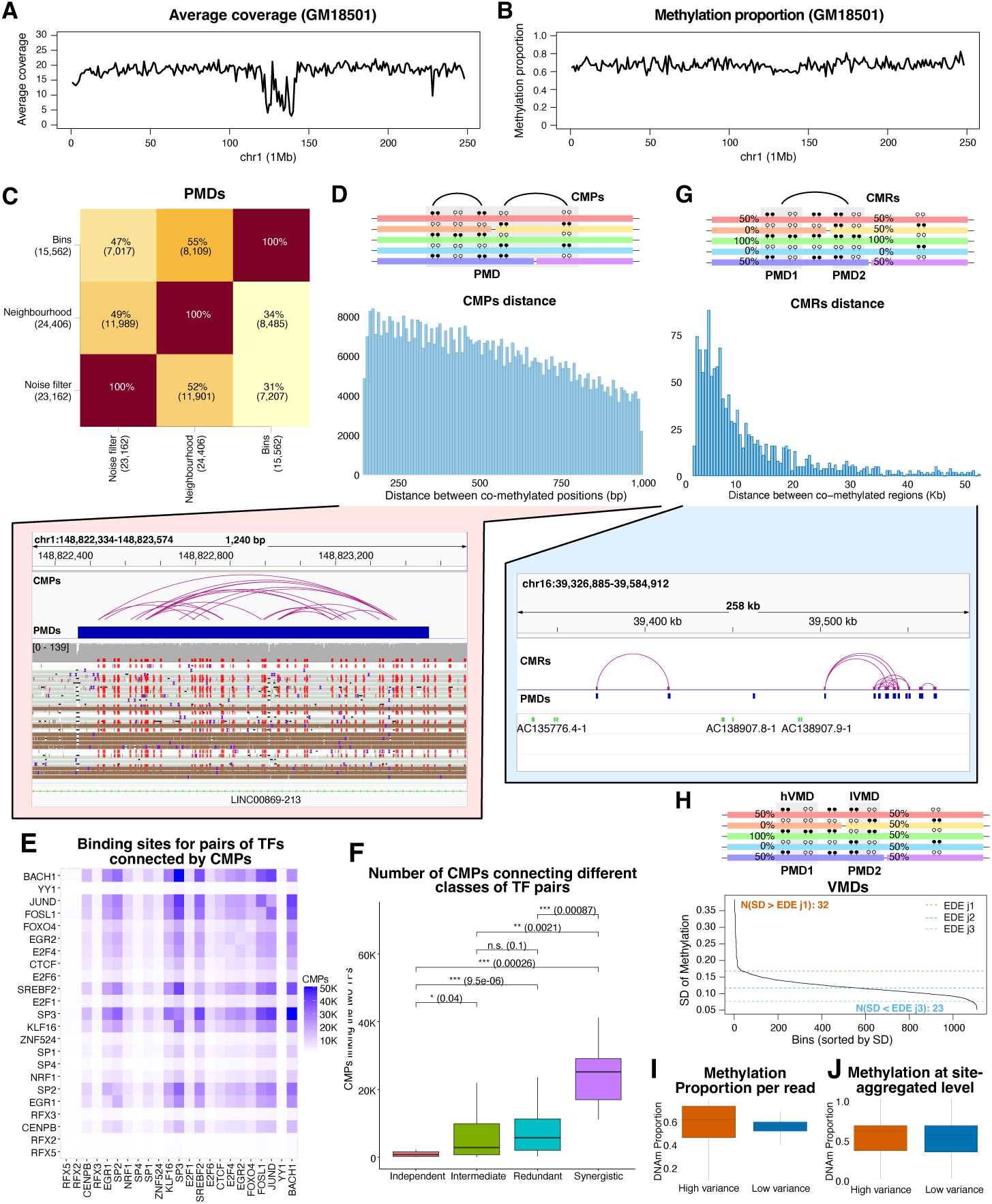
Single-molecule methylation profiling and heterogeneity analysis in the GM18501 lymphoblastoid cell line. (*A-B*) Low-resolution profile for coverage and average methylation across chr1: (*A*) mean sequencing coverage and (*B*) mean methylation proportion. The x-axis represents the position on chr1, while the y-axis the average coverage or methylation proportion. We consider 1 Mb tilling bins and computed the average coverage or methylation proportion in those bins. (*C*) Detection of Partially Methylated Domains (PMDs) with methylation proportion between 0.4 and 0.6 using three different methods: bins, neighbourhood and noise filter. The heatmap shows the overlap between the PMDs detected with the method on the x-xis to the ones detected with the method on the y-axis. For the downstream analysis we used PMDs detected with bins method. (*D*) Distribution of distance between comethylated sites within PMDs. Top panel shows a cartoon representing CMPs. Bottom panel shows an IGV browser visualisation of co-methylation structure within a representative 2 *Kb* PMD in *LINC00869-213* on chromosome 1. (*E*) Number of CMPs connecting binding sites for different TF pairs. Darker colours indicate that there are more CMPs linking the binding sites of the two TFs. (*F*) Comparison of the number of CMPs connecting binding sites for different TF pairs for different classes of predicted TF cooperative interactions in HepG2. We performed a Mann-Whitney U test and used Bonferroni correction for multiple testing. (*G*) Distribution of distance between co-methylated regions computed over PMDs. Top panel shows a cartoon representing CMRs. Bottom panel shows an IGV browser visualisation of co-methylation structure between PMDs on chr16. (*H*) Distribution of weighted standard deviation (SD) of per-read methylation proportions across 1,109 PMDs. Dashed lines indicate the *ede*-derived thresholds separating highly variable (hVMDs; SD *>* 0.185, *n* = 32) and low-variable (lVMDs; SD *<* 0.090, *n* = 23) methylated domains. To define high- and low-variability groups, we applied the *ede* algorithm from the R package *inflection*, which detects knee points in the standard deviation distribution. Top panel shows a cartoon representing VMDs. (*I-J*) Box plots comparing methylation proportion distributions between a hVMD (chr1:127,131,967-127,133,166) and a lVMD (chr1:129,959,267-129,961,366) at the: (*I*) single-read level and (*J*) site-aggregated level. At cytosine aggregate level there is no difference in variation of methylation proportions, but at read level hVMD shows higher variability than lVMD.

#### Detection of Partially Methylated Domains

Non-coding regions of the DNA called enhancers control the activity of genes by determining whether a gene is transcribed or not (36; 37) and recent work has shown that intermediary levels of DNA methylation characterises active enhancers (38; 39). This indicates that enhancers would be located in Partially Methylated Domains (PMDs), i.e., extended genomic regions characterised by intermediate methylation levels that reflect large-scale epigenetic heterogeneity. Thus, identification of PMDs would be useful to identify potential regulatory regions of the genome. Because definitions of partial methylation vary across studies (12; 13; 40), here, we selected to defined PMDs as domains with mean methylation proportions between 0.4 and 0.6. Given that fully methylated or unmethylated regions typically show proportions close to 1 or 0, respectively, restricting average methylation within PMDs to the [0.4, 0.6] interval allows us to specifically capture domains with balanced mixtures of methylated and unmethylated CpGs. Given that the 5mC levels display correlation between cytosines up to 2*Kb* (Figure S2B), we decided to select this size for the bins/windows when detecting PMDs.

Next, we used the three methods newly developed to detect PMDs that are based on previously implemented methods for DMRs detection, namely: noise-filter (using a smoothing kernel on the total number of reads and methylated reads), neighbourhood (considering individual Cs and merging them iteratively), and bins (tilling the genome in fix size bins and computing average methylation per bin) (16). Genomic overlap among the resulting PMD sets was used to assess detection consistency. The bins method displayed the lowest number of PMDs ( 15*K*) and approximately half of those were detected by the two other methods (Figure 3C). For the downstream analysis, we decided to use the PMDs detected by bins method, but the results from the other methods can be used as well.

#### Detection of co-methylated CpGs

Transcription Factors (TFs) are protein that bind to DNA at specific sites within promoters or enhancers to control transcription of target genes (41). Many TFs do bind cooperatively (42) to control gene expression and several previous studies have reported that the binding of a large percentage of TFs seem to be affected by DNA methylation (43; 44; 45; 46). One possibility is that, at binding sites of TFs that display DNA methylation sensitivity and bind cooperatively, CpGs would display co-methylation patterns. Single molecule analysis of DNA methylation allows us to investigate patterns of co-methylation between CpGs.

To take advantage of this, we developed a new method for identifying co-methylation positions (CMPs) called `computeCoMethylatedPositions`, which quantifies if co-occurrence of methylation at two different CpGs is more than expected by chance (Figure 3D). In regions that are fully methylated or unmethylated, it is not possible to assess co-methylation between CpGs and, thus, we detected these CMPs within PMDs. As expected, the total number of CMPs gradually decreased with distance (Figure 3D). Figure 3D also shows an example of co-methylation patterns on chromosome 1 at a PMD located in LINC00869-213 locus.

#### Prediction of TF co-binding from CMPs

We hypothesised that CMPs could also be used to identify co-binding of TFs with DNA methylation sensitive binding. Thus, we analysed the binding sites of 305 human TFs that contain a CpG in their motif and are expressed in LCL cells (see *Materials and Methods*). We considered CMPs that connect two CpGs located within 150 bp (nucleosome size) of each other and found that 27 TFs showed motif enrichment at these CMPs. For some of these TFs, there is a large number of CMPs that link their predicted binding sites (see Figure 3E), indicating that these TFs could display co-binding.

To further investigate if there is additional evidence supporting the co-binding of these TFs, we considered a classification of TF co-operative interactions in HepG2 cell line from (47). In particular, TF pairs were classified into four groups: independent, redundant, intermediate or synergistic; with the latter displaying the highest evidence for cooperative interaction and the former the lowest. 131 of the TF pairs we identified in our CMPs analysis were classified in HepG2 into one of these four categories; see Figure S3A. Synergistic TF pairs show the highest number of CMPs linking their binding sites, while independent pairs show the lowest number, with intermediate levels for the intermediate and redundant groups; see Figure 3F. This indicates that for TF pairs that were predicted to bind cooperatively there is a high number of CMPs linking their binding sites, while for the ones that were predicted to bind independently the number of CMPs is low. Interestingly, 6 of the 7 synergistic TF pairs involve at least one subunit of AP-1 (JUND-FOSL1, JUND-JUND, FOSL1-FOSL1, JUND-BACH1, JUND-KLF16 and FOSL1-SP2), with the last one including CTCF (CTCF-SREBF2) a TF known for having its binding impacted by DNAm (48). There is a strong effect of DNAm on binding to DNA for JUND and FOSL1 (both part of AP-1 complex) and SREBF2 (45), while there is lower effect of DNAm on KLF16 and SP2 binding to DNA. Altogether, this indicates that the CMPs can be used to identify synergistic co-binding of TFs and distinguish that from independent cobinding.

#### Detection of co-methylated regions

Transcription of a gene is controlled by the activity of a promoter and in many cases of multiple enhancers. One possibility to detect all potential regulatory regions is to identify which regions display co-occuring epigenetic states, such as co-methylation. To detect co-methylation of regions, we considered the average proportion methylation within two regions at single molecule level. If methylation level is highly correlated (when one region is highly methylated the other one is also highly methylated and vice versa), this would indicate that the two regions are potentially co-regulated or co-regulate the same gene; Figure 3E. Similar as in the case of CMPs, for regions that are fully methylated or unmethylated, it is not possible to assess co-methylation between them and, thus, we detected co-methylation of PMDs. We detected CMRs in GM18501 cells using the `computeCoMethylatedRegions` function. Given the average length of ONT reads of approximately 30 *Kb*, we considered PMDs within 50 *Kb* of each other and observed that the closer two PMDs are the more likely it is they are co-methylated (Figure 3G). Interestingly, we detect that most co-methylated regions are within 20 *Kb* of each other, but we also detect CMRs with distances of up to 50 *Kb*. The fact that most CMRs detected are within 20 *Kb* of each other could also be affected by the average length of ONT read being 30 *Kb*, which results in fewer PMDs at longer distances being covered by multiple reads in order to be able to detect CMRs. It is worthwhile noting that enhancers within 50 *Kb* distance of target genes can regulate in cis without the need of 3D contacts (49). Figure 3G also shows an examples of co-methylated regions (PMDs) on chromosome 16.

#### Detection of Variable Methylated Domains

DNA methylation variability is high within a population and while at least half is associated with genetic variation (50), some has been linked with environment exposure (51). Most studies so far has investigated DNA methylation variability between CpGs within a sample or between individuals. Nevertheless, having DNA methylation data at single molecule levels allows to detect regions that display high variability within a sample. We developed a new function called `computeVMDs` that allows detection of variable methylated domains (VMDs). In particular, `computeVMDs` computes the average percentage methylation within a region on each read and then estimates the variability of that average methylation over all reads. For this analysis, we used the PMDs identified previously as potential VMDs. In total, we identified 32 highly variable methylated domains (hVMDs; SD *>* 0.185) and 23 lowly variable methylated domains (lVMDs; SD *<* 0.090) among 1,109 PMDs (Figure 3H), i.e. the PMDs with sufficient per-read coverage (`minReadsPerCytosine` = 4, `minCytosinesCount` = 4) to estimate read-level variance.

Having classified VMDs into high- and low-variability groups, we examined the variance in methylation proportion for one hVMD and lVMD at both the site-aggregated (at individual CpG) and single-read levels. Interestingly, the two regions considered (the hVMD and the lVMD) are non-coding with the nearest gene (LOC124905671) being over 1 *Mb* away. At the single-read level, hVMDs regions exhibited high difference in variance compared to lVMDs (Fligner–Killeen test, *P* = 3.89 *×* 10*^−^*^5^) despite a similar median methylation proportion (Mann-Whitney U test, *P* = 0.553); Figure 3I. In contrast, at the site-aggregated level, hVMDs and lVMDs showed similar methylation proportions and variability (Mann-Whitney U test, *P* = 0.531; Fligner–Killeen test, *P* = 0.659) (Figure 3J). Overall, this shows that using single molecule DNA methylation we can identify new epigenetically variable regions of the genome that could have not been detected using bulk approaches (such as WGBS or EPIC array).

### Characterisation of genome wide sex differences in DNA methylation in lymphoblastoid cell lines

Next, we applied these new methods to investigate sex differences in DNA methylation. For this purpose, we selected two lymphoblastoid cell lines (LCLs) from the 1000 Genomes ONT dataset (17): HG00277 (male) and HG00326 (female). To characterise global methylation patterns prior to differential analysis, we first generated low-resolution profiles based on 1 Mb windows aligned to the T2T-CHM13 reference genome. On chr1, both cell lines showed consistent sequencing coverage of approximately 10 reads across most regions, with total read counts (ReadsN) at CpG sites showing broadly comparable distributions between the two cell lines (Figure 4A and Figure S4A). Whilst the overall methylation proportions were broadly similar between the two cell lines, the female cell line exhibited a marginally higher methylation proportion and correspondingly elevated methylated read counts than the male on chromosome 1 (Figure 4B). This supports our previous findings in a large cohort where we found that majority of sex associated differentially methylated positions on autosomes were hypermethylated in females compared to males (24).

**Figure 4:**
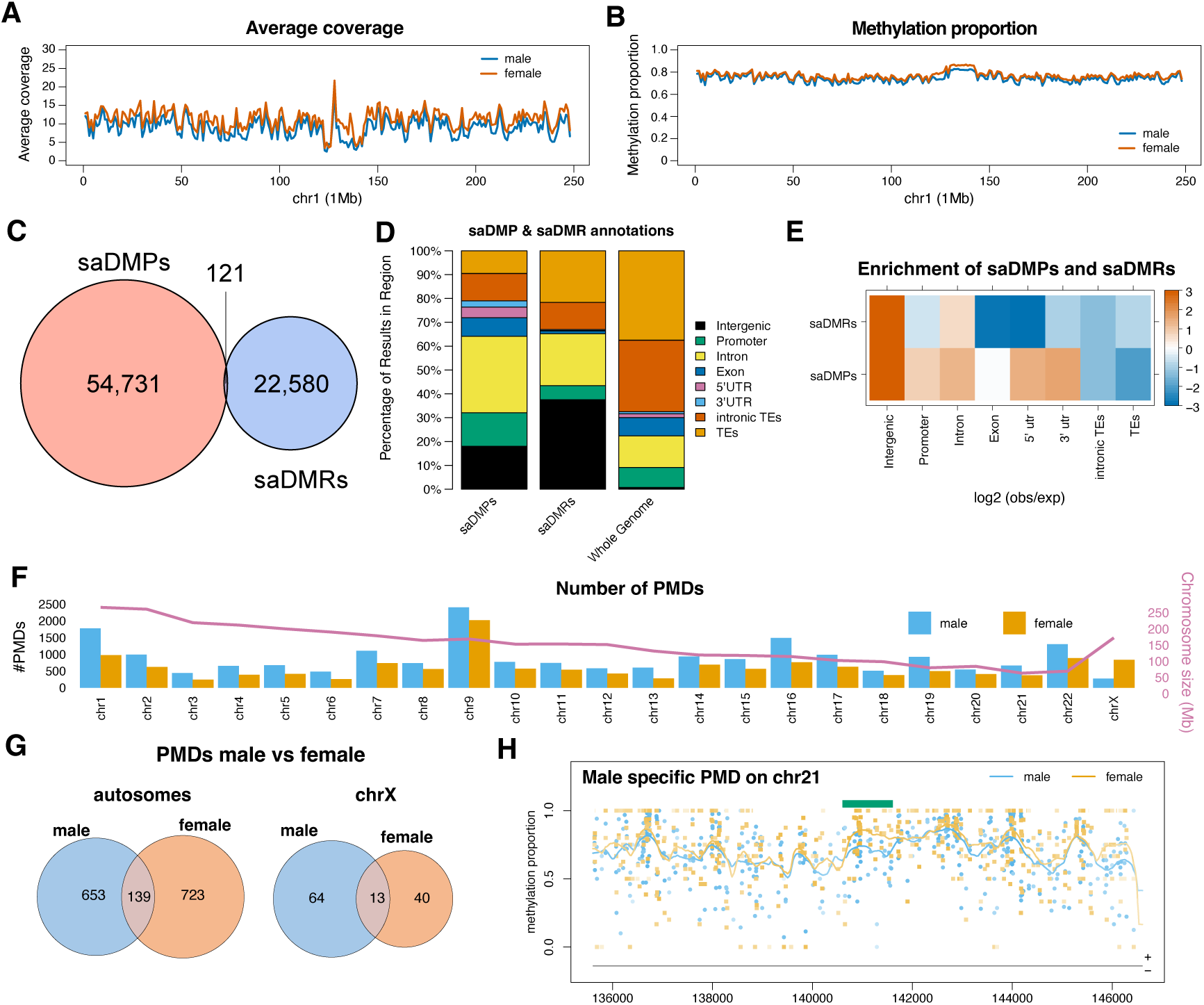
Investigation of sex-associated DNAm differences in lymphoblastoid cell lines. (*A-B*) Low-resolution profiles for: (*A*) coverage and (*B*) average DNAm levels of HG00277 (male) and HG00326 (female) across chr1. The x-axis represents the position on chr1, while the y-axis the average coverage or methylation proportion. We consider 1 Mb tilling bins and computed the average coverage or methylation proportion in those bins. (*C*) Genome-wide overlap between saDMRs (bin size 400 bp, minimum proportion difference 0.4) in the two LCL cell lines and saDMPs identified with EPIC array on a large cohort (24). (*D-E*) Annotation of saDMRs and saDMPs relative to genomic features. (*E*) shows the *log*_2_(*obs/exp*) based on the whole genome distribution of the different annotations, with orange showing enrichment and blue depletion. (*F*) Number of PMDs per chromosome in male and female samples. The magenta line represents the chromosome size. (*G*) Overlap of PMDs that have at least 20 reads in both males and females, except on chrX in males where we used a threshold of 10 reads (in order to account that there is only one copy of chrX in males). We considered the case of autosomes and chrX separately. (*H*) An example of one of the male specific PMDs on autosomes (chr21). The points on the graph represent methylation proportion of individual cytosines, their colour (sky blue or orange) which sample they belong to and the intensity of the colour how many reads that particular cytosine had (darker colours indicate more reads, while lighter colours indicate fewer reads). The solid lines represent the smoothed profiles and the intensity of the colour the coverage at the corresponding position. The location of the PMD is marked by the green rectangle.

To systematically investigate sex associated differences on autosomes, we first computed DMRs between the male and female LCL lines (saDMRs) and we compared these to the sex-associated differentially methylated positions (saDMPs) reported in (24). Figure 4C shows a low overlap between saDMRs and saDMPs, which is not surprising given that in (24) we analysed whole blood samples and here we are analysed LCL lines, thus, reflecting biological differences between systems. One reason for the low overlap is that the EPIC array covers only 3% of CpGs in the human genome, while ONT can analyse all CpGs. The saDMRs are specifically enriched in intergenic and intronic regions, while the saDMPs are mostly enriched in promoters, 5’UTR and 3’UTR (Figure 4D-E), which reflects the bias in the probes on the EPIC array. This can explain why some regions are detected only in the ONT data and are not detected on the EPIC, but does not explain why the majority of saDMPs detected in the EPIC data are not discovered in the ONT data. One possibility is that, in (24), we analysed approximately 3.6*K* samples which ensured the statistical power to detect very small but statistically significant differences (e.g., of 1%), which is not possible here with only 2 samples. For the saDMRs, we did use a threshold of 0.4 in methylation and using a lower threshold can detect more, but still not the majority of saDMPs (Figure S4B). Finally, another possibility is the non homogenous coverage of the genome in the ONT data affects the calling of DMRs. Indeed, we found a lower coverage in the male cell line over the saDMPs specifically with only 20% of the saDMPs from the epic study having 5 reads or more (Figure S4C), which confirms why some of these regions were not detected from the ONT data.

To investigate co-methylation structure within partially methylated domains, we identified PMDs in both cell lines. Using the previous more stringent thresholds or larger domain sizes resulted in a very low number of PMDs and, thus, for this analysis, we used a methylation proportion range of [0.2, 0.8] with a minimum domain size of 1 *Kb*. Our results showed that males have a slightly higher number of PMDs on autosomes and females have higher number of PMDs on chrX; Figure 4F. One possible explanations the fact that X inactivation in females leads to a higher number of intermediary levels in methylation of chromosome X (52; 53). Another possibility is that some of these differences in PMDs between males and females could be caused by differences in local sequencing depth. To correct for this, we selected all PMDs that have on average at least 20 reads in both males and females, except on chrX in males where we used a threshold of 10 reads (in order to account that there is only one copy of chrX in males). Interestingly, this resulted only between 16% and 18% of PMDs detected in both males and females on autosomes, supporting previous findings that there are large differences in DNA methylation on autosomes between sexes (24); Figure 4G. Figure 4H shows one example of a male specific PMD located on chr21 that is located in a non-coding region ( 30 *Kb* downstream of LOC124908168), where there is a drop in DNA methylation in males compared to females.

### Co-methylation of positions and regions in the male and female LCL cell lines

Next, we investigated the presence of co-methylation positions (CMPs) in males and females. As expected, in male specific PMDs, the majority of identified CMPs are male specific, while, in female specific PMDs, the majority of identified CMPs are female specific; Figure S5A-B. Thus, we turned our attention to CMPs in the 152 PMDs (Figure 4G) that are shared between males and females. Our results showed that only 512 CpG pairs show co-methylation in both males and females, while a similar number of pairs show co-methylation in males only Figure 5A. Interestingly, the number of co-methylated CpG pairs specific to females is more that double compared to the ones specific to males. None of the CMPs are located at imprinted loci; Figure 5A and Figure S5A-B. Further dissecting the 512 common CMPs between males and females, we found that only 30% display similarly high statistical significance, while 21% show a bias towards males and a similar percentage a bias towards females; Figure 5B. Figure 5C shows an example of female/male specific CMPs on chromosome 5.

**Figure 5:**
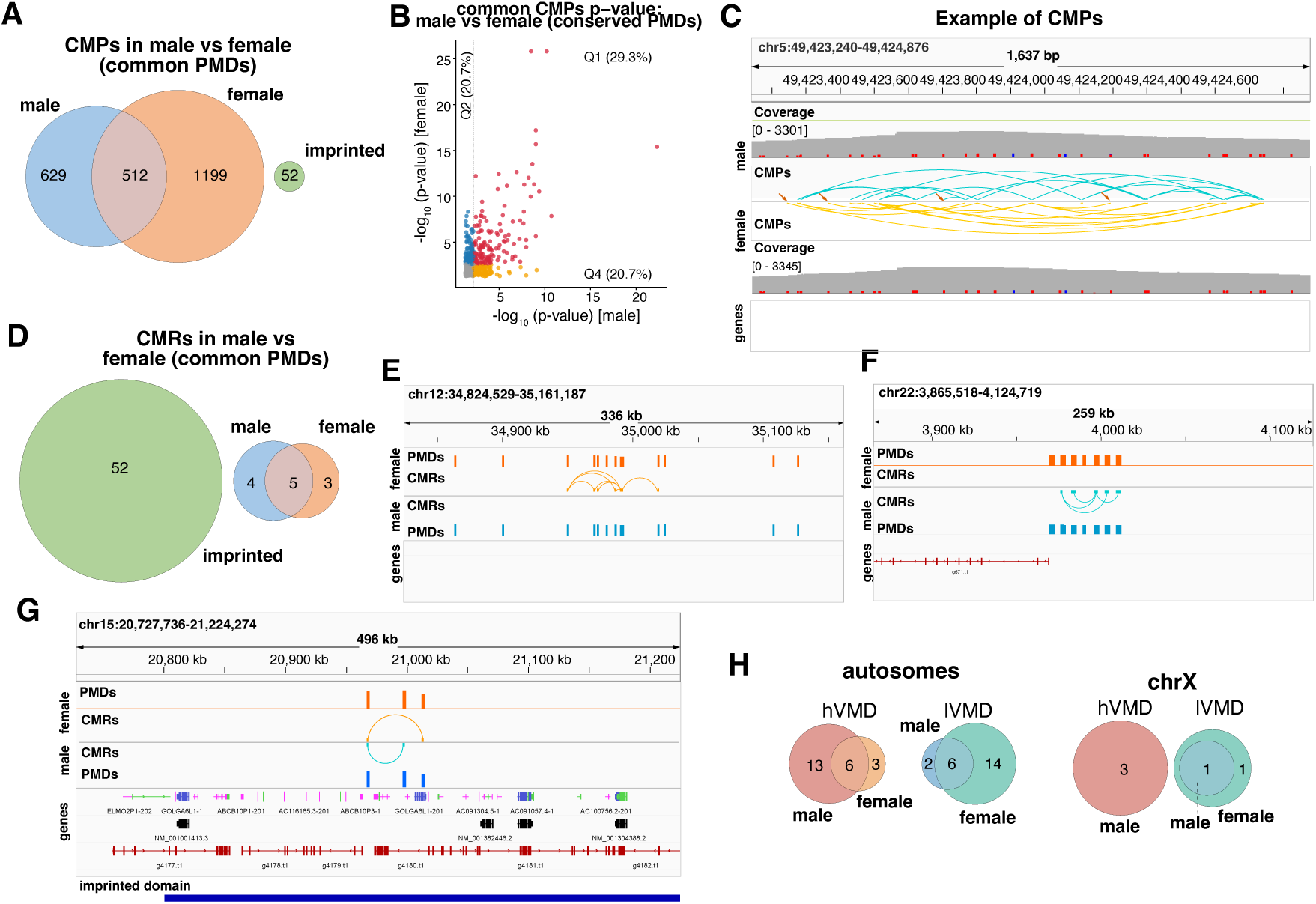
Co-methylation differences between males and females in lymphoblastoid cell lines. (*A*) The common and sex specific CMPs located in common PMDs and their relationship with imprinted loci. (*B*) Comparison between the p-values of the common CMPs between males and females. (*C*) IGV browser visualisation of female/male specific CMPs at a locus (chr5:49,423,240-49,424,876). Orange arrows highlight some of the sex specific CMPs. (*D*) The common and sex specific CMRs considering only the common PMDs and their relationship with imprinted loci. (*E*) IGV browser visualisation of female specific CMRs at locus (chr12:34,824,529-35,161,187). The arcs show the two CpGs that are co-methylated and we represent male CMPs by blue colour and female ones by yellow. (*F*) IGV browser visualisation of male specific CMRs at locus (chr22:3,865,518-4,124,719). The arcs show the two regions that are co-methylated and we represent male CMRs by blue colour and female ones by orange.(*G*) IGV browser visualisation of a CMRs at the imprinted locus (chr15:20,727,736-21,224,274). (*H*) Overlap between hVMDs and lVMDs in male and female samples. We considered the case of autosome and chrX separately.

Next, we characterised co-methylated regions (CMRs) and assess whether correlated methylation patterns could be detected and whether those are specific to males or females. To compare the CMRs between males and females, we considered only regions with consistent coverage in both. For PMDs that are present in both males and females, we do identify 4 CMRs that are male specific and 3 that are female specific, with 5 being present in both males and females. These low numbers are influenced by the small number of PMDs common between males and females that have at least 20 reads and are within 50 *Kb* of each other. A higher sequencing depth would likely result in higher number of both CMPs and CMRs. Figure 5E-F show the female and male specific CMRs. The male specific CMR (Figure 5F) is located near the gene *g671.t1*, whereas the female specific CMR (Figure 5E) is not located near any gene.

Similarly as for CMPs, in male specific PMDs we identified mainly male specific CMRs, while in female specific PMDs mainly female specific CMRs; Figure S5C-D. Interestingly, one of the CMRs between male specific PMDs is at an imprinted locus (chr15:20,727,736-21,224,274). This CMR is near GOLGA6L26 gene and is plotted in Figure 5G.

#### Variable methylated domains in the male and female LCL cell lines

Next, we evaluated the new function to detect VMRs filterVMRsONT between males and females. To ensure that these VMRs are not driven by differences in methylation, we considered only regions with consistent coverage in both samples. Thus, we used the common 152 PMDs between males and females (Figure 4G) and all promoters (2 *Kb* upstream of the TSS), which resulted in no VMR being detected. One possibility is that we had a very small number of regions to test if they act as VMRs and thus we also considered all promoters (2 *Kb* upstream of TSS) and this did not identify any VMR either. This suggest that there is no difference in variability per read between males and females in the samples we considered. This could be either due to this variation between males and females not being present in LCL cells or due to the coverage of the samples not being high enough to detect these differences.

An alternative approach is to detect VMDs in each sample and then compare the overlap of hVMDs or lVMDs between the samples. Figure 5H confirms that in the 152 common PMDs between male and female samples, there are only 22 VMDs in males and 9 in females, with only 6 being common between the two. These low numbers further confirm the lack of detected VMRs. All sex-specific hVMDs are non-coding, with the 3 female specific hVMDs being at least 715 *Kb* away from nearest gene. Nevertheless, there are 4 male specific hVMDs that are within 250 Kb of three genes, namely: (*i*) FRG1FP.(chr22:12,834,990-12,836,089, 109 *Kb* away), (*ii*) ANKRD30BP2 (chr21:11,227,422-11,228,421, 154 *Kb* away and chr21:11,166,422-11,166,921, 215 *Kb* away) and (*iii*) LOC124908437 (chr22:2,519,840-2,520,939, 197 *Kb* away). Interesting, ANKRD30BP2 has 2 hVMDs within 250 *Kb* that are male specific and which is aberrantly expressed in 58% of prostate tumours (54)

#### Identification of 5hmC and non-CpG methylation

ONT long read sequencing allows estimation of both methylation and hydroxymethylation in CpG context or non-CpG context. To investigate whether these epigenetic modifications can be detected and analysed by DMRcaller, we used Dorado v0.9.6 with the `sup,5mC_5hmC` model to extract base modification calls in six cytosine contexts. Our results showed that while 5mC at CpG sites is on average around 0.6-0.8 genome wide (Figure 3B and Figure 4B), hydroxymethylation or non-CpG methylation is significantly lower; see Figure S6. Only 5hmC in CG context is slightly above 10% with the rest being below that (between 2% and 6%). Given that this particular sample had on average 10 reads per position, this indicates that these samples do not have the required sequencing depth to detect such low level of methylation.

## Discussion

Here we present a new framework that support native Oxford Nanopore Technologies (ONT) sequencing data, which is integrated within the *DMRcaller* R/Bioconductor package. The `readONTbam` function enables direct import of base modification calls from ONT-aligned BAM files, supporting a range of cytosine modification contexts including CpG 5mC, 5hmC, and non-CpG CHG and CHH methylation. In particular, the framework supports identification of molecule-to-molecule heterogeneity in DNA methylation (VMDs and VMRs), and coordinated methylation (CMPs and CMRs). Together, these capabilities provide a unified framework for characterising the full landscape of epigenetic variation from ONT data, spanning population-level differential methylation, read-level heterogeneity, and long-range co-methylation structure, within a single, accessible Bioconductor package. By unifying VMD identification, CMR analysis, and DMR calling within a single Bioconductor package, *DMRcaller* moves beyond the bulk-level aggregation and accuracy-focused benchmarking that characterise most existing ONT methylation tools(55; 5; 56), enabling integrated characterisation of methylation heterogeneity at both population and single-molecule resolution within a single analytical framework.

It is worthwhile noting that DNA methylation data is inferred from ONT long reads using machine learning models to predict when a base is methylated or not-methylated. This means that DNAm is not a directly detected by sequencing and, as a result, there might be differences between base calling tools or versions of the same tools (57). *DMRcaller* relies on the input from the bam file generated by Dorado, which also offers the possibility that in the future, that newer models could display better accuracy at estimating DNAm.

A key finding from the single-sample analysis was that high-variance methylated domains (hVMDs) and low-variance methylated domains (lVMDs) are indistinguishable at the site-aggregated level, with comparable median methylation proportions (0.60 vs 0.553), IQRs, and variances, and no significant difference in distribution or variance (Mann-Whitney U test, *P* = 0.531; Fligner–Killeen test, *P* = 0.659). In contrast, at the single-read level, hVMDs exhibited an 8.48-fold greater variance and a 3.45-fold wider IQR than lVMDs, with a highly significant difference in dispersion (Fligner–Killeen test, *P* = 3.89 *×* 10*^−^*^5^). This demonstrates that read-level methylation heterogeneity, rather than a shift in mean methylation proportion, is the distinction that is entirely invisible to bulk-level analyses. This observation is consistent with recent reports showing that single-molecule ONT data reveal substantial methylation heterogeneity within PMDs that is masked by population-averaged metrics (7), and highlights the value of the single-molecule resolution afforded by ONT sequencing over array- or bisulfite-based approaches that necessarily report population-averaged methylation levels(58; 56).

Furthermore, we show how CMPs linking CpGs within 150 bp of each other can be used to detect co-binding of TFs. Interestingly, we predict elevated co-binding for several pairs of TFs with some of these being previously reported to display synergistic co-binding (47). The majority of pairs of TFs predicted to have co-binding involved AP-1 subunits, including homodimers (JUND and FOSL1) and heterodimers (JUND-FOSL1) (59). This provides evidence that some of the predicted co-binding events by our approach are indeed cooperative binding.

Application of the new functionality to a male–female lymphoblastoid cell line pair from the 1000 Genomes ONT dataset (17) identified sex-associated DMRs (saDMRs) that partially overlapped with the large-cohort saDMP resource reported in (24). The latter was derived from EPIC array data encompassing over 3,600 individuals. Whilst 121 genome-wide overlaps were confirmed, the genomic distributions of saDMRs and saDMPs were markedly divergent: saDMRs were predominantly intergenic (0.375) and intronic (0.218), whereas saDMPs were enriched in introns, promoters, and UTRs, which are regions well represented on the EPIC array but not necessarily captured at higher density by unbiased ONT sequencing. This divergence most likely reflects the complementary nature of the two platforms: the EPIC array targets a predefined set of approximately 850,000 CpGs with known regulatory relevance and is powered by large sample numbers, whilst ONT sequencing provides unbiased coverage of approximately 27 million CpG sites genome-wide (roughly 32-fold more), making it better suited to detecting methylation variation in repetitive and intergenic regions that are systematically under-represented on arrays(56; 60; 58). In addition, we also found a significantly reduced coverage in the male LCL cell line of the saDMPs identified on the EPIC array. This highlights that one limitation of the technology is the non-homogenous coverage of the genome. Nevertheless, this could be addressed by increasing the overall sequencing depth to ensure that majority of the genome will have enough coverage to detect these differences.

Co-methylation analysis is only possible with long read sequencing technology. Array do not allow capturing this co-methylation structures, while short read sequencing could potentially only capture co-methylated positions up to 150 *bp*. Nevertheless, that is within one nucleosome length and would miss interesting biological mechanisms that span over multiple nucleosomes. Furthermore, being able to identify co-methylated regions within up to 50 *Kb* of each other would open to possibility to detect cisregulatory regions. In particular, we know that enhancers within 50 *Kb* distance of target genes can regulate in cis without the need of 3D contacts (49) and our method could detect these candidate enhancers or co-regulated promoters.

Co-methylation analysis within PMDs revealed a higher number of female CMPs at PMDs that are common between males and females. Nevertheless, we detected only a small number of CMRs in the male and female sample (less than 10). Given that we detected a significantly higher number of CMRs in the GM18501 cell line that had a higher coverage suggests that our method is dependent of sequencing depth. Furthermore, we found that computing differences in PMDs, VMDs, CMPs and CMRs between samples is robust only at regions with consistent medium or high coverage (minimum of at least 10 reads, but 20 reads would be recommended).

## Conclusions

In this work, we present a new computational framework integrated within *DMRcaller* package that enables integrated genome-wide methylation analysis from native ONT sequencing data at both population and single-molecule resolution. By introducing new functions for VMR detection, VMD identification, CMP and CMR analysis alongside the existing DMR framework, we provide a unified analytical platform that captures epigenetic variation across multiple levels of resolution within a single, accessible workflow. Together, these results establish *DMRcaller* as a flexible and scalable framework for single-molecule epigenomic profiling from long-read sequencing data, with broad applicability to studies of epigenetic heterogeneity, differential methylation, and co-methylation structure across diverse biological contexts.

## Author’s contributions

Y.J.K. and N.R.Z. conceived and designed the experiments. Y.J.K. performed the experiments and analysed the data. N.R.Z. supervised the work. Y.J.K. and N.R.Z. wrote the paper.

## Acknowledgements

We thank members of the Centre for Epigenetics at QMUL and Zabet lab members (especially Archie Simpson) for discussion and feedback on this manuscript. The analysis was performed on the High Performance Computing Facility at the Queen Mary University of London.

