## Supplementary Material for "A Framework for Quantifying DNA Methylation Heterogeneity and Detecting Co-methylated loci from Native Nanopore Sequencing"

immediate

### Supplementary Figures

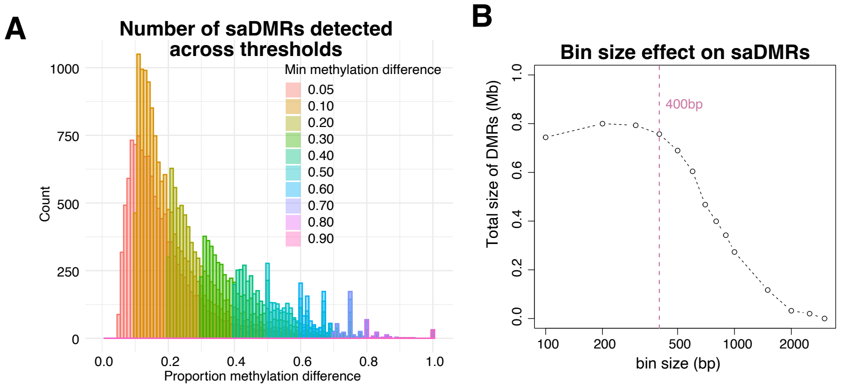


Figure S1: *Detecting sex-associated DMRs in lymphoblastoid cell lines.* (*A*) Histogram of saDMRs for different methylation proportion difference. We considered different thresholds for the methylation proportion differences between 0.05 and 0.9. (*B*) Total size of DMRs (in Mb) as a function of bin size used to detect DMRs. For bins larger than 400 bp, the total size of regions called as DMRs starts reducing. DMR were called in CpG context using the bins method with the following parameters: `test` = `"score"`, `pValueThreshold` = 0.05, `minCytosinesCount` = 1, `minGap` = 200 bp, `minSize` = 50 bp, and `minReadsPerCytosine` = 2.

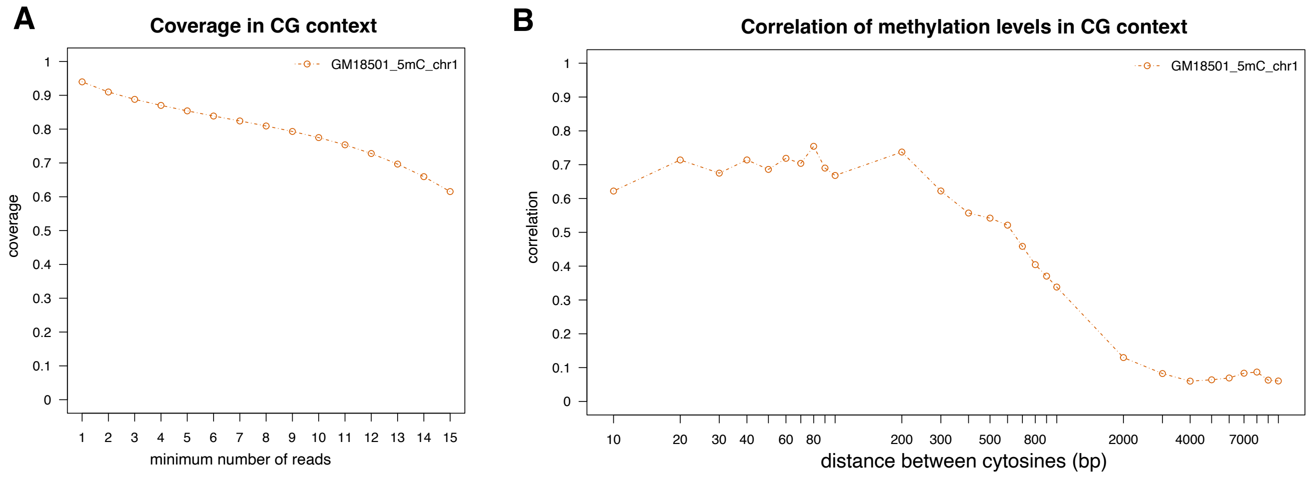


Figure S2: *Analysis in the GM18501 lymphoblastoid cell line.* (*A*) Coverage of the ONT DNAm data in CpG context. The graph plots the proportion of CpGs that have at least the number of reads specified on the x-axis. (*B*) Spatial correlation between DNAm at CpG sites in the genome. CpGs within distances of 200 bp show high correlation in DNA methylation level, which corresponds to size of the nucleosome and linker DNA. For distances of 2 Kb or larger, there is negligible correlation between methylation levels of the CpG sites.

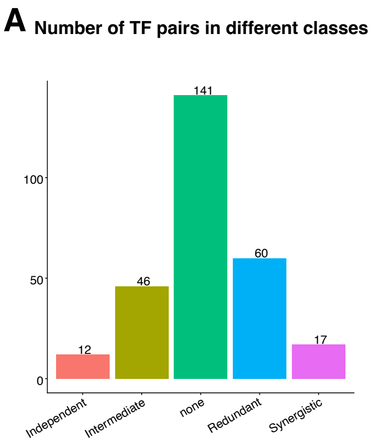


Figure S3: *TF co-binding prediction in GM18501 cell line from CMP data.* (*A*) Number of TF pairs considered in our analysis that were classified as Independent, Intermediate, Redundant or Synergistic in HepG2 cells. None represents the TF pairs that were not classified in the HepG2 cell line.

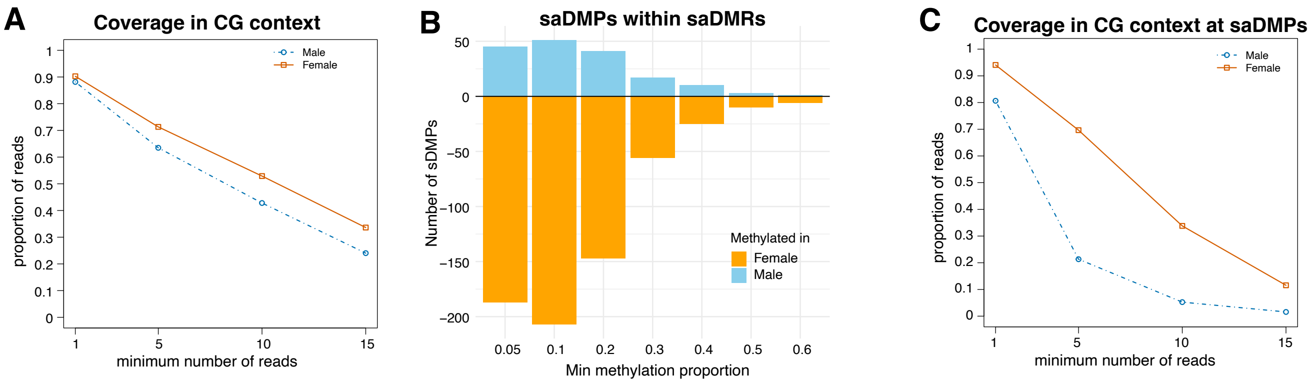


Figure S4: *Characterising sex-associated DNAm differences in lymphoblastoid cell lines.* (*A*) Coverage of the ONT DNAm data for 5mC in the two LCL cell lines. The graph plots the proportion of CpGs that have at least the number of reads specified on the x-axis. (*B*) Number of saDMRs overlapping saDMPs reported in a large cohort study using EPIC array. We considered different levels for the min proportion difference thresholds of 0.05–0.6 and performed the analysis on chromosome 1. The hypermethylated saDMRs in female are presented in orange and saDMRs hypermethylated in males are represented in blue. The number of saDMPs on chr1 that were hypermethylated in females was 3,558 and hypermethylated in males was 2,064. (*C*) Coverage of the ONT DNAm data for 5mC in the two cell lines at saDMPs only.

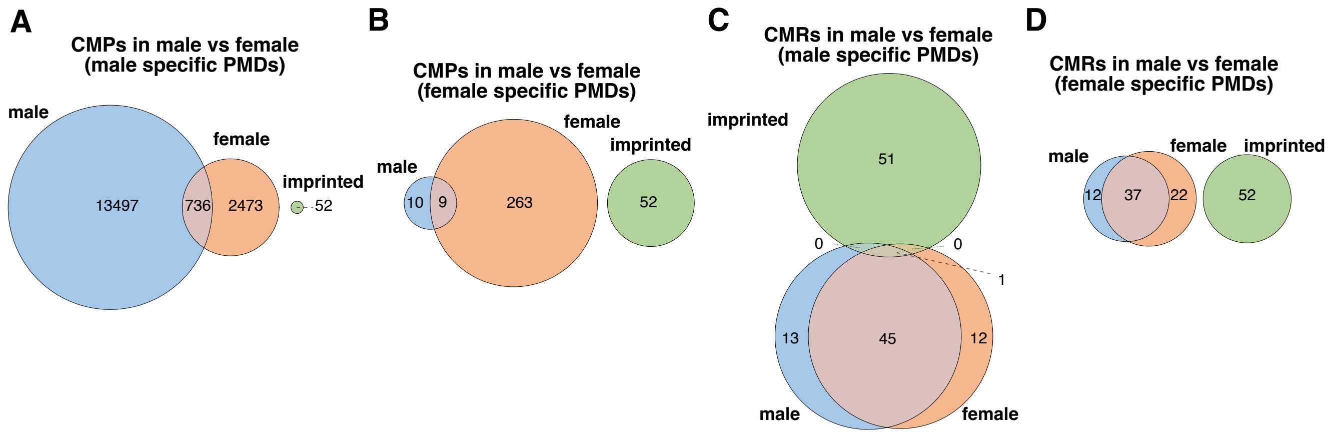


Figure S5: *Co-methylation differences between males and females in lymphoblastoid cell lines.* (*A*-*B*) The overlap of male and female specific CMPs that were detected in male specific PMDs (*A*) or female specific PMDs (*B*) and their relationship with imprinted loci. (*C*-*D*) The overlap of male and female specific CMRs that were detected in male specific PMDs (*C*) or female specific PMDs (*D*) and their relationship with imprinted loci.

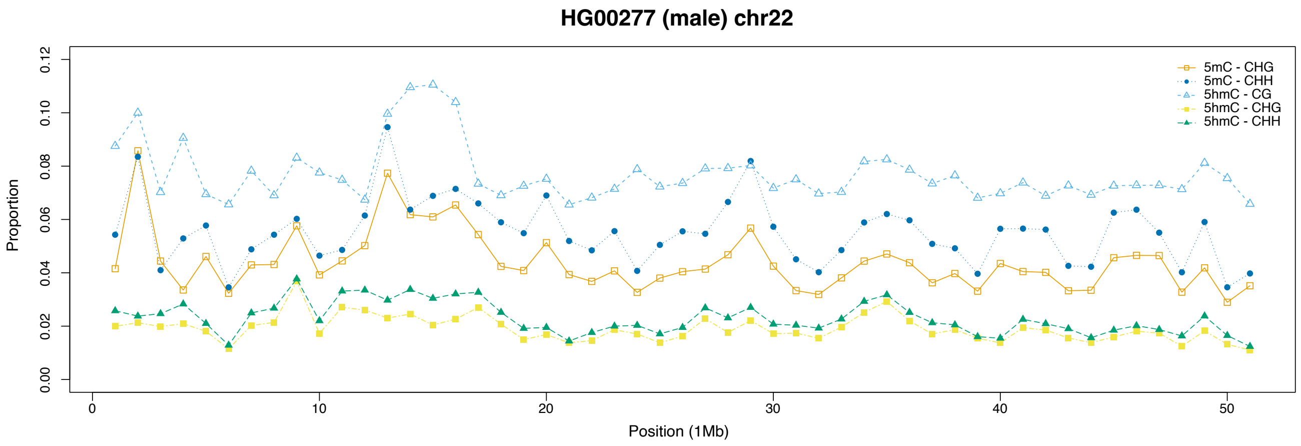


Figure S6: *Detection of 5hmC and non-CpG methylation.* Low-resolution profiles for average 5mC and 5hmC levels in CpG and non-CpG contexts of HG00277 (male) across chr22. The x-axis represents the position on chr22, while the y-axis the average methylation proportion. We consider 1 Mb tilling bins and computed the average methylation proportion in those bins. 5mC in non-CpG and 5hmC display very low levels of DNA methylation, with only 5hmC displaying approximately 10% methylation on average.
